## Supplementary information for "The capsule and genetic background, rather than specific individual loci, strongly influence *in vitro* pneumococcal growth kinetics"

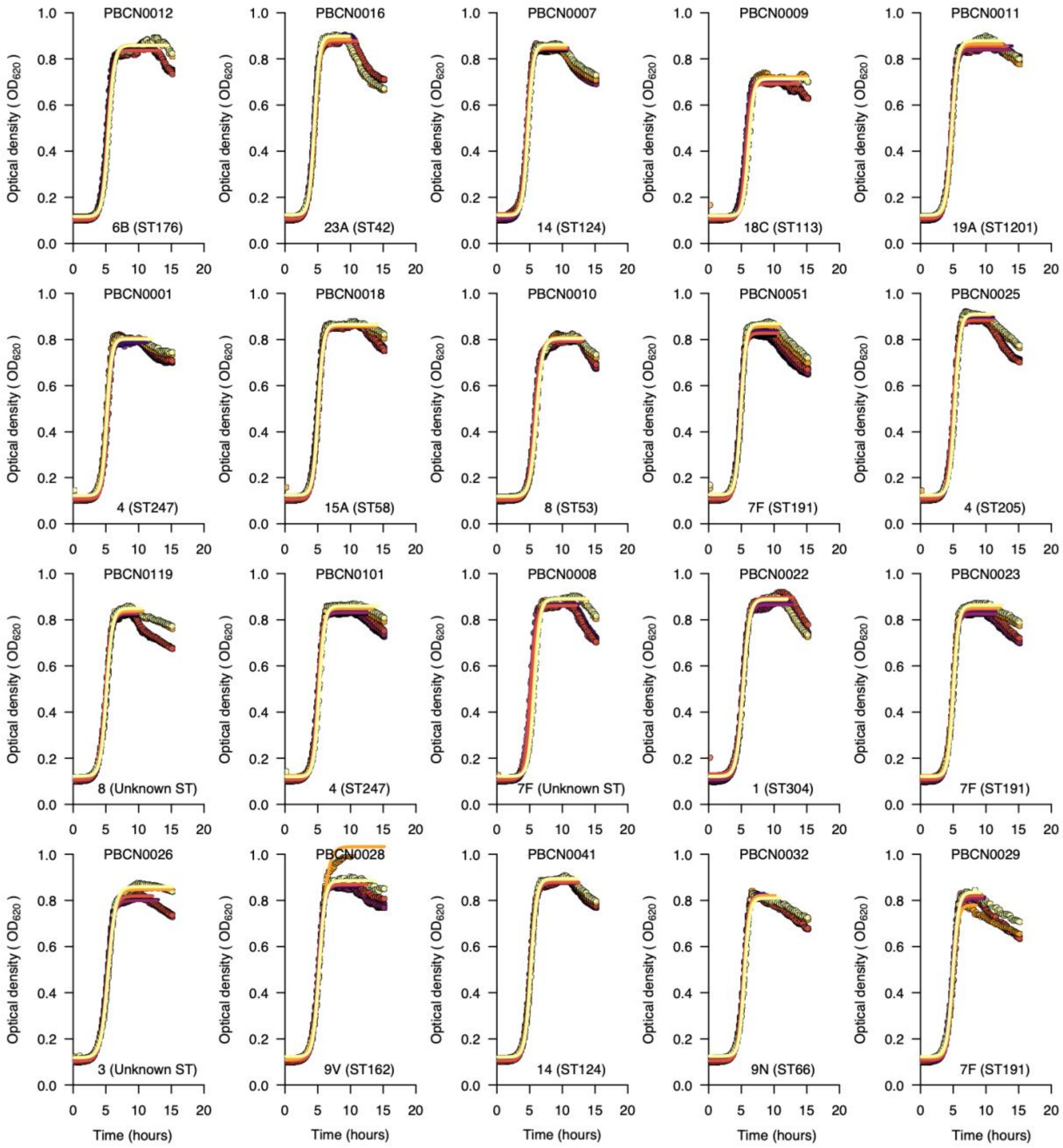


**Supplementary Fig. 1. Growth curves for selected serotypes show the *in vitro* pneumococcal growth dynamics and estimated growth features inferred by fitting a statistical model.** The growth curves above show the optical density at 620nm monitored over 15 hours for an example dataset of 20 pneumococcal isolates representing different serotypes and sequence types (STs) based on the pneumococcal MLST scheme. We used the nonlinear optimisation algorithm to identify a combination of parameters with the best fit to a logistic curve based on the lowest residual sum of squares for the predicted value based on the model relative to the observed growth data (see methods). We fitted separate curves based on a subset of points for each replicate up to the end of the stationary phase or end of the experiment, whichever came first. We did this to get a better fit for the logistic curve during the growth phase, as it does not allow for negative growth, which happened after the stationary phase. We fitted the curves to infer the growth features for each of the six replicates. The averaged estimates for each replicate were then used for the downstream analyses.

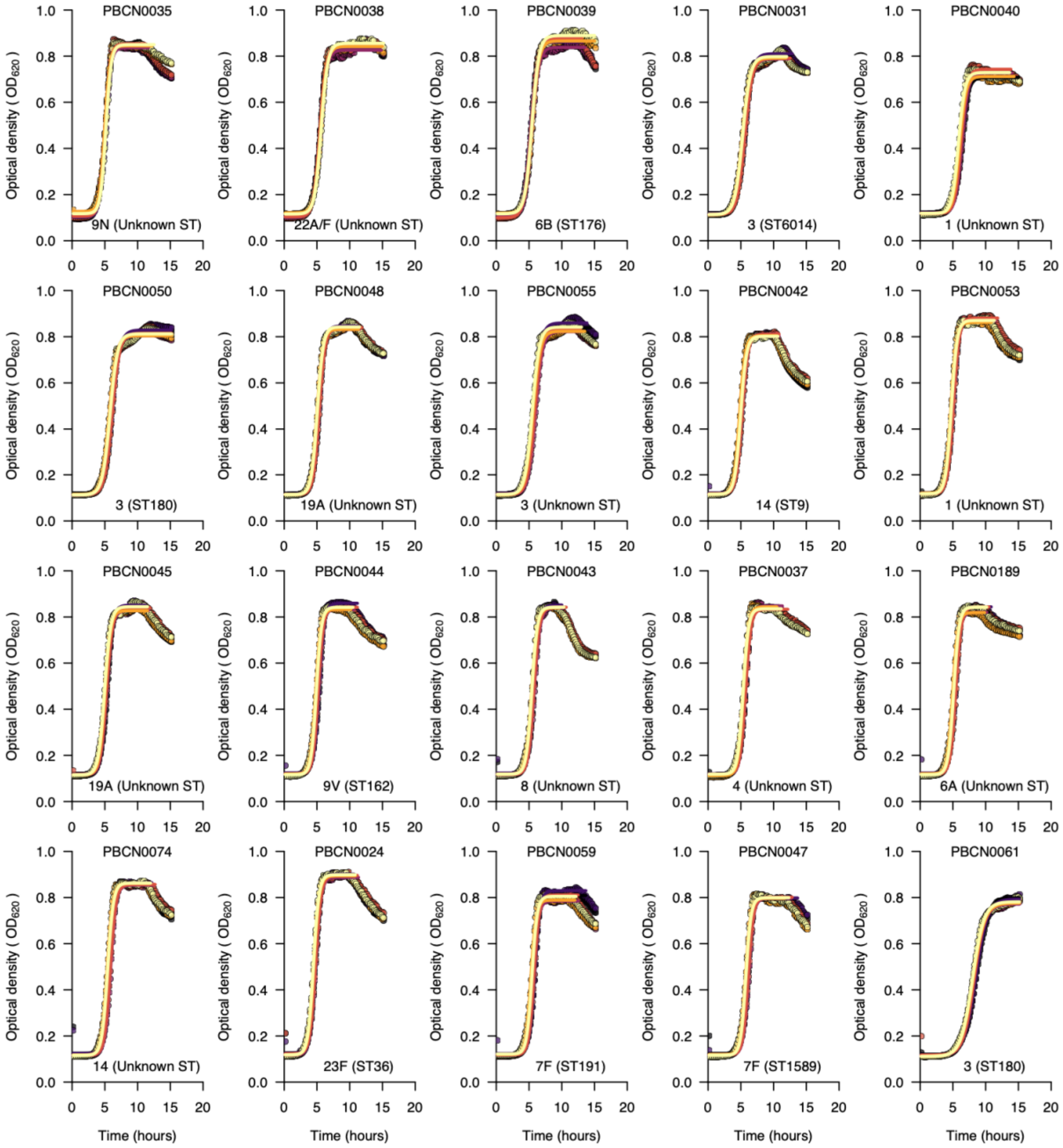


**Supplementary Fig. 1 (continued). Growth curves for selected serotypes show the *in vitro* pneumococcal growth dynamics and estimated growth features inferred by fitting a statistical model.** The growth curves above show the optical density at 620nm monitored over 15 hours for an example dataset of 20 pneumococcal isolates representing different serotypes and sequence types (STs) based on the pneumococcal MLST scheme. We used the nonlinear optimisation algorithm to identify a combination of parameters with the best fit to a logistic curve based on the lowest residual sum of squares for the predicted value based on the model relative to the observed growth data (see methods). We fitted separate curves based on a subset of points for each replicate up to the end of the stationary phase or end of the experiment, whichever came first. We did this to get a better fit for the logistic curve during the growth phase, as it does not allow for negative growth, which happened after the stationary phase. We fitted the curves to infer the growth features for each of the six replicates. The averaged estimates for each replicate were then used for the downstream analyses.

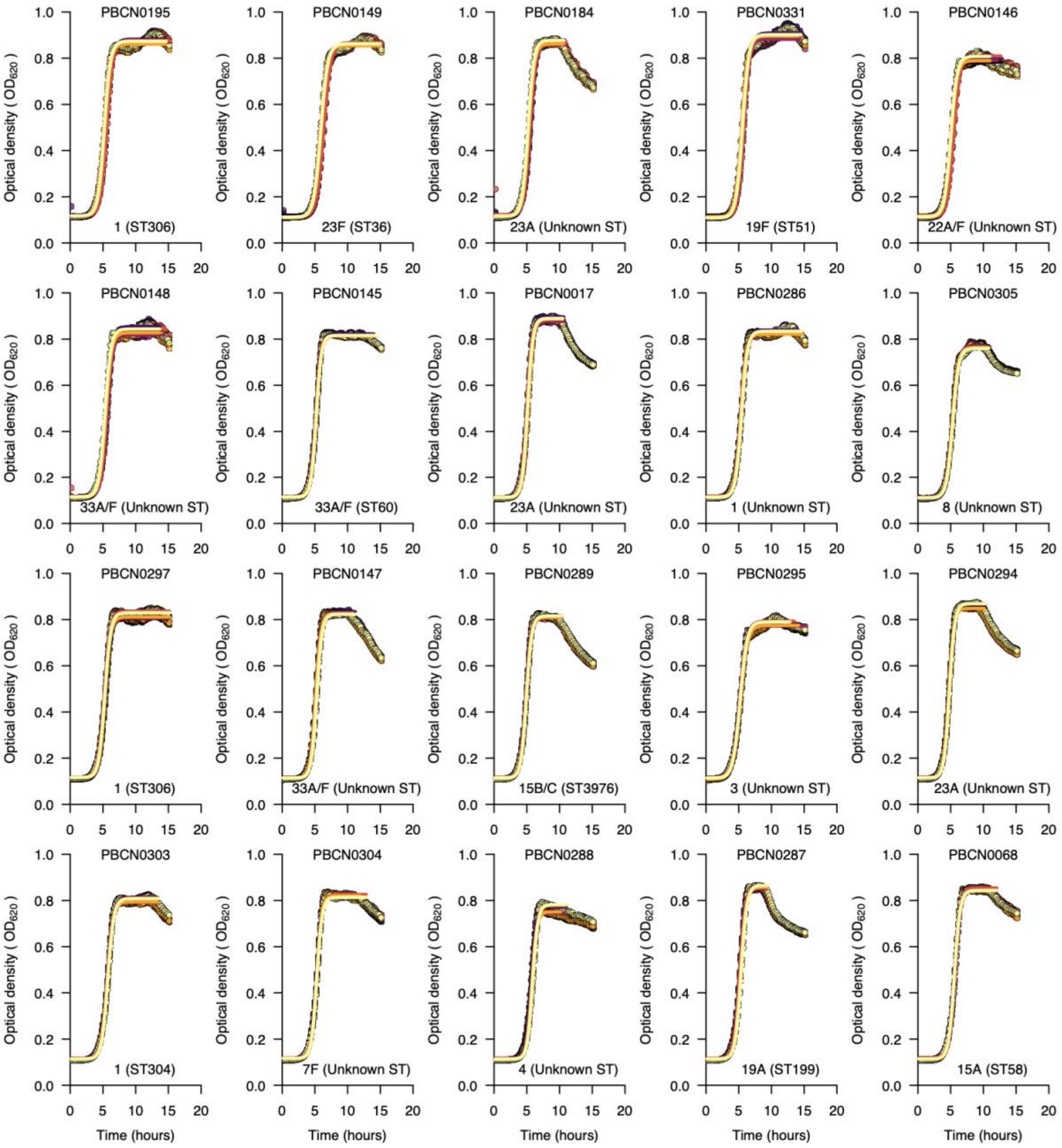


**Supplementary Fig. 1 (continued). Growth curves for selected serotypes show the *in vitro* pneumococcal growth dynamics and estimated growth features inferred by fitting a statistical model.** The growth curves above show the optical density at 620nm monitored over 15 hours for an example dataset of 20 pneumococcal isolates representing different serotypes and sequence types (STs) based on the pneumococcal MLST scheme. We used the nonlinear optimisation algorithm to identify a combination of parameters with the best fit to a logistic curve based on the lowest residual sum of squares for the predicted value based on the model relative to the observed growth data (see methods). We fitted separate curves based on a subset of points for each replicate up to the end of the stationary phase or end of the experiment, whichever came first. We did this to get a better fit for the logistic curve during the growth phase, as it does not allow for negative growth, which happened after the stationary phase. We fitted the curves to infer the growth features for each of the six replicates. The averaged estimates for each replicate were then used for the downstream analyses.

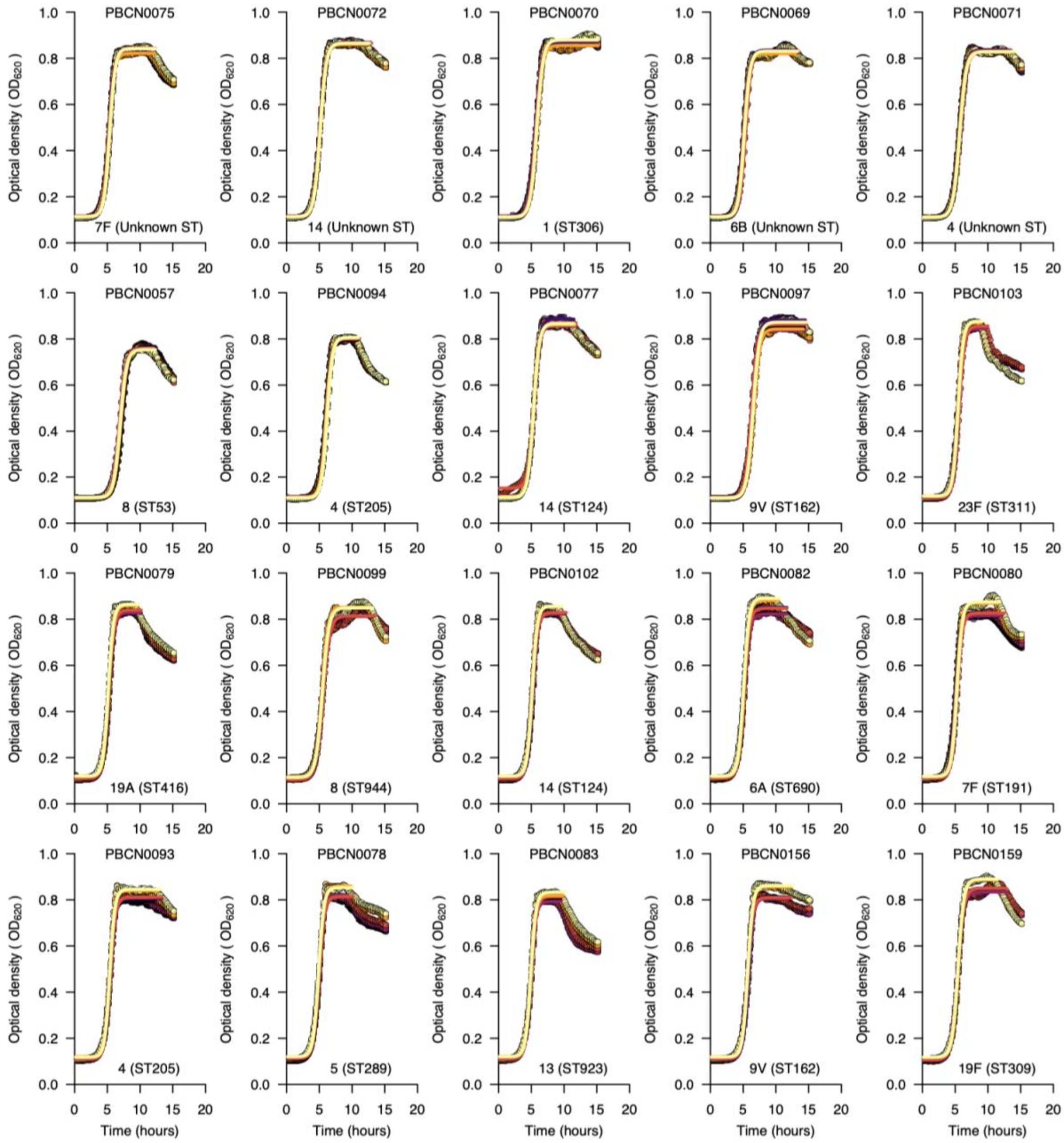


**Supplementary Fig. 1 (continued). Growth curves for selected serotypes show the *in vitro* pneumococcal growth dynamics and estimated growth features inferred by fitting a statistical model.** The growth curves above show the optical density at 620nm monitored over 15 hours for an example dataset of 20 pneumococcal isolates representing different serotypes and sequence types (STs) based on the pneumococcal MLST scheme. We used the nonlinear optimisation algorithm to identify a combination of parameters with the best fit to a logistic curve based on the lowest residual sum of squares for the predicted value based on the model relative to the observed growth data (see methods). We fitted separate curves based on a subset of points for each replicate up to the end of the stationary phase or end of the experiment, whichever came first. We did this to get a better fit for the logistic curve during the growth phase, as it does not allow for negative growth, which happened after the stationary phase. We fitted the curves to infer the growth features for each of the six replicates. The averaged estimates for each replicate were then used for the downstream analyses.

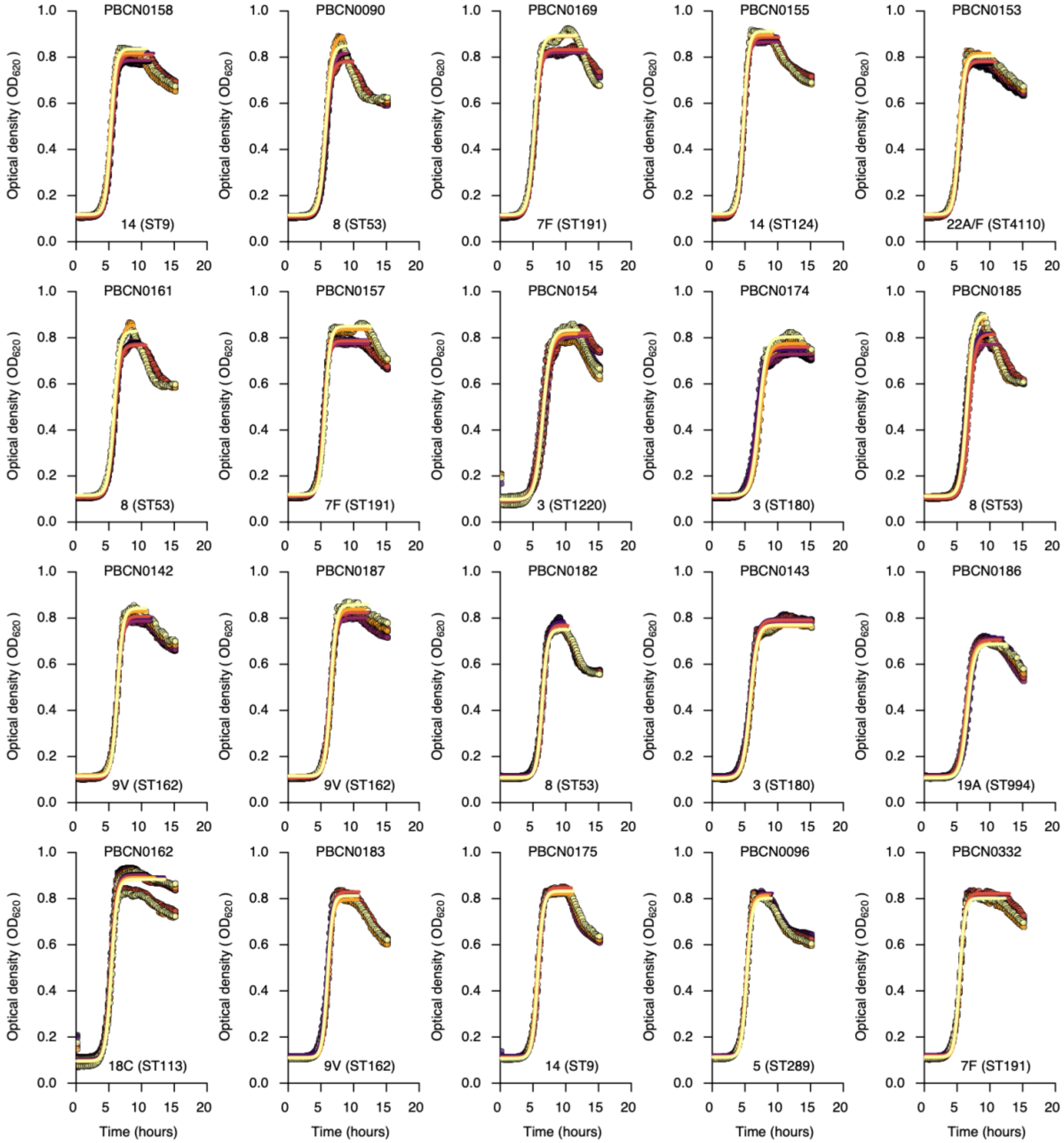


**Supplementary Fig. 1 (continued). Growth curves for selected serotypes show the *in vitro* pneumococcal growth dynamics and estimated growth features inferred by fitting a statistical model.** The growth curves above show the optical density at 620nm monitored over 15 hours for an example dataset of 20 pneumococcal isolates representing different serotypes and sequence types (STs) based on the pneumococcal MLST scheme. We used the nonlinear optimisation algorithm to identify a combination of parameters with the best fit to a logistic curve based on the lowest residual sum of squares for the predicted value based on the model relative to the observed growth data (see methods). We fitted separate curves based on a subset of points for each replicate up to the end of the stationary phase or end of the experiment, whichever came first. We did this to get a better fit for the logistic curve during the growth phase, as it does not allow for negative growth, which happened after the stationary phase. We fitted the curves to infer the growth features for each of the six replicates. The averaged estimates for each replicate were then used for the downstream analyses.

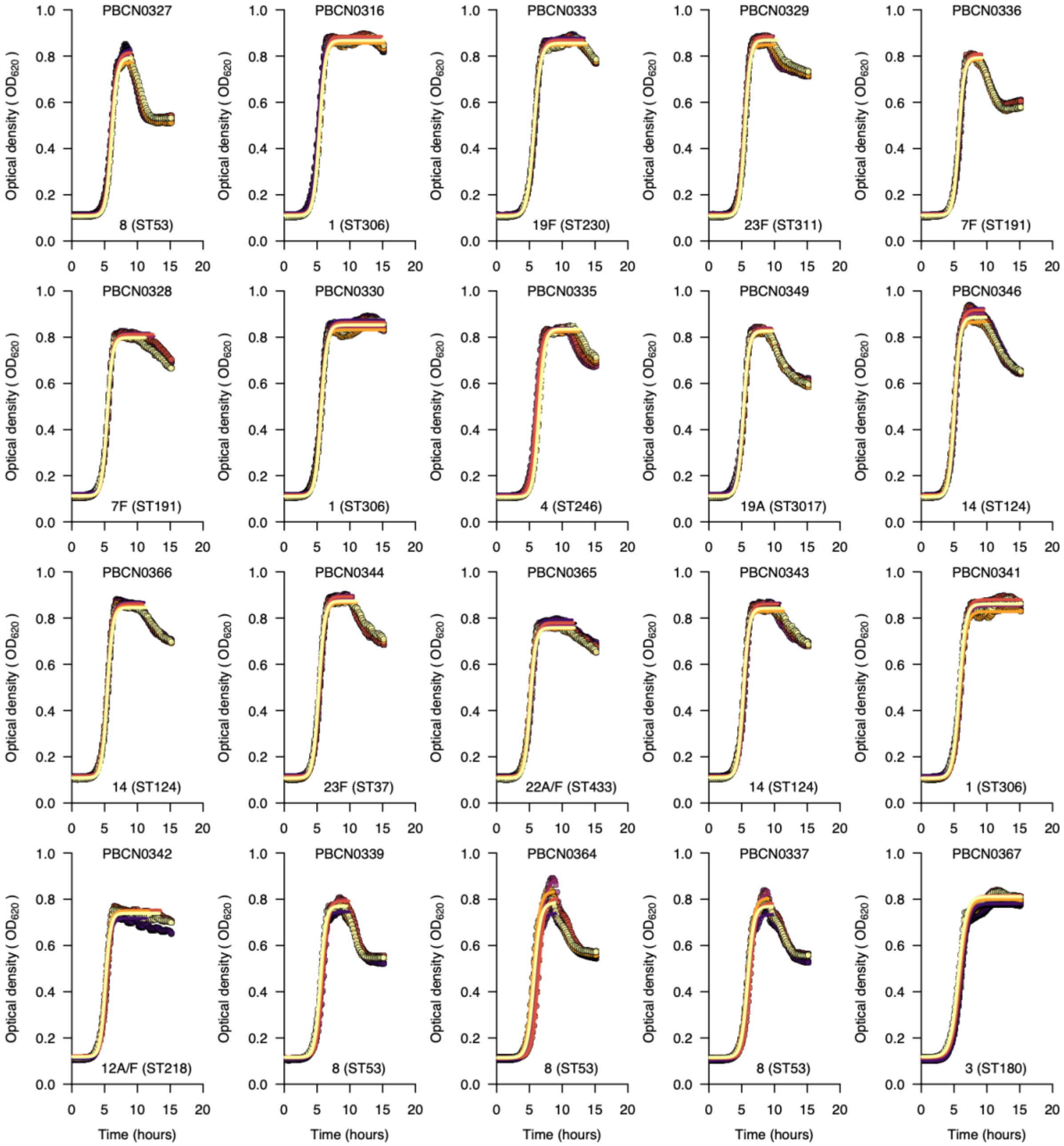


**Supplementary Fig. 1 (continued). Growth curves for selected serotypes show the *in vitro* pneumococcal growth dynamics and estimated growth features inferred by fitting a statistical model.** The growth curves above show the optical density at 620nm monitored over 15 hours for an example dataset of 20 pneumococcal isolates representing different serotypes and sequence types (STs) based on the pneumococcal MLST scheme. We used the nonlinear optimisation algorithm to identify a combination of parameters with the best fit to a logistic curve based on the lowest residual sum of squares for the predicted value based on the model relative to the observed growth data (see methods). We fitted separate curves based on a subset of points for each replicate up to the end of the stationary phase or end of the experiment, whichever came first. We did this to get a better fit for the logistic curve during the growth phase, as it does not allow for negative growth, which happened after the stationary phase. We fitted the curves to infer the growth features for each of the six replicates. The averaged estimates for each replicate were then used for the downstream analyses.

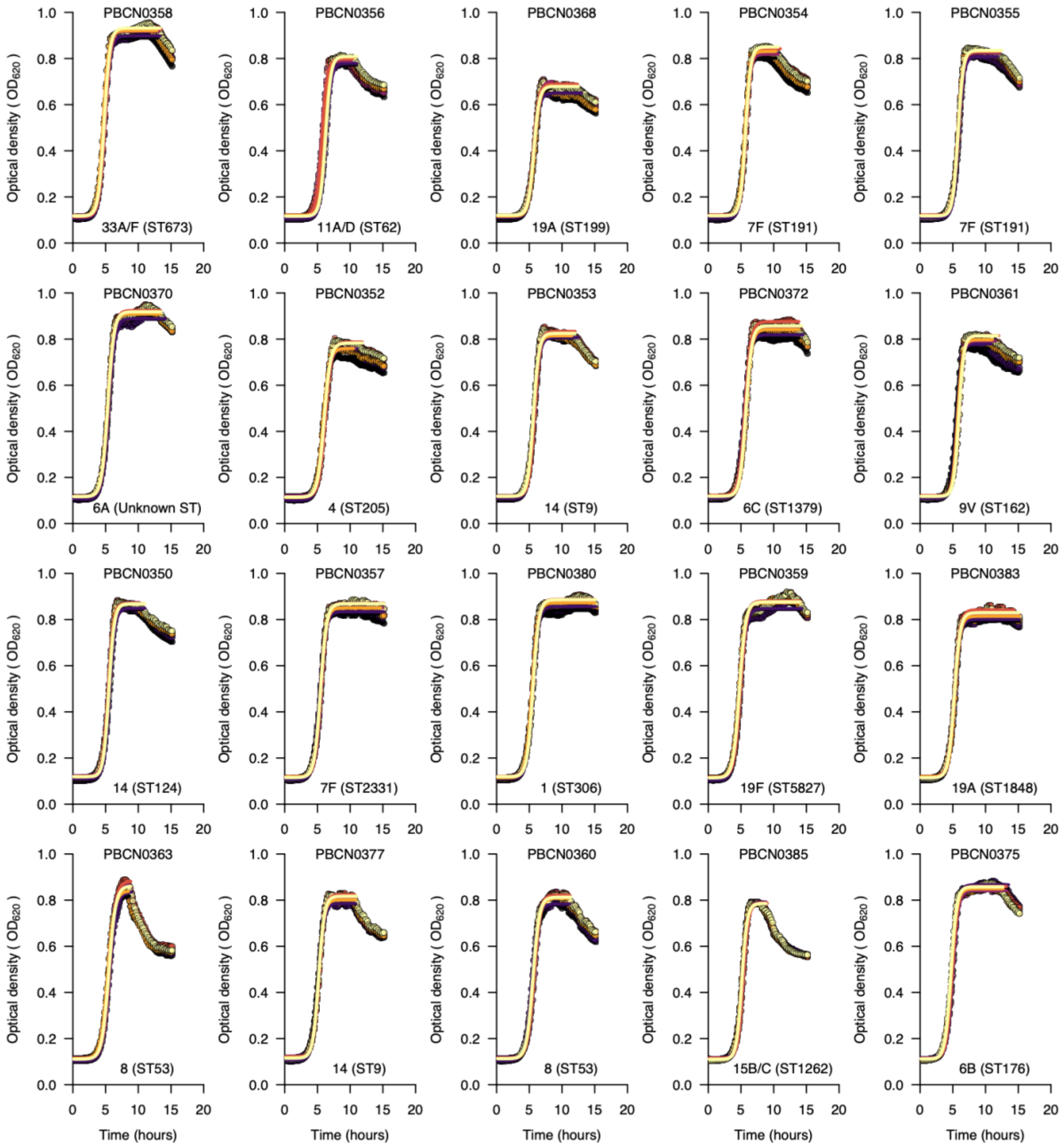


**Supplementary Fig. 1 (continued). Growth curves for selected serotypes show the *in vitro* pneumococcal growth dynamics and estimated growth features inferred by fitting a statistical model.** The growth curves above show the optical density at 620nm monitored over 15 hours for an example dataset of 20 pneumococcal isolates representing different serotypes and sequence types (STs) based on the pneumococcal MLST scheme. We used the nonlinear optimisation algorithm to identify a combination of parameters with the best fit to a logistic curve based on the lowest residual sum of squares for the predicted value based on the model relative to the observed growth data (see methods). We fitted separate curves based on a subset of points for each replicate up to the end of the stationary phase or end of the experiment, whichever came first. We did this to get a better fit for the logistic curve during the growth phase, as it does not allow for negative growth, which happened after the stationary phase. We fitted the curves to infer the growth features for each of the six replicates. The averaged estimates for each replicate were then used for the downstream analyses.

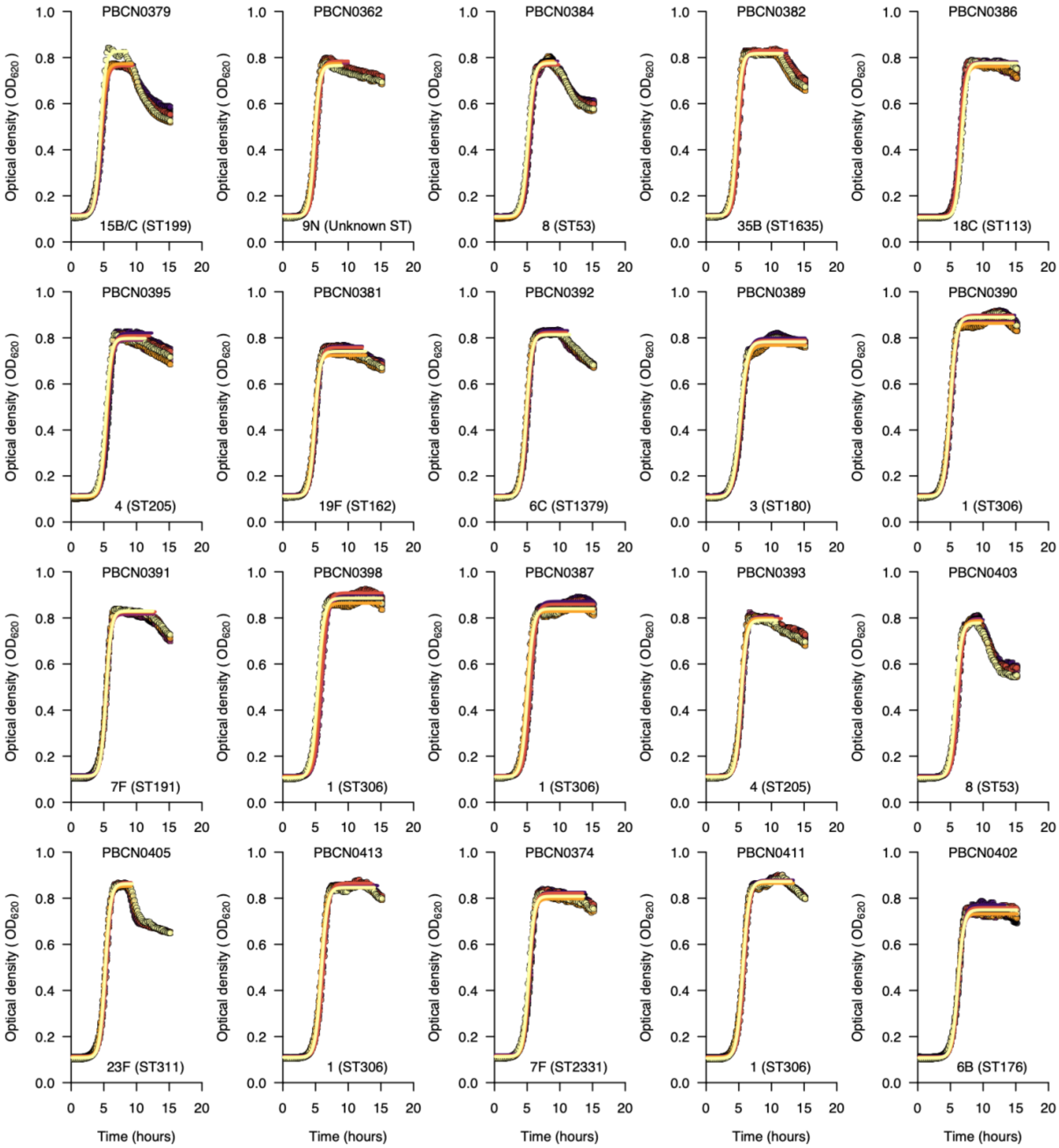


**Supplementary Fig. 1 (continued). Growth curves for selected serotypes show the *in vitro* pneumococcal growth dynamics and estimated growth features inferred by fitting a statistical model.** The growth curves above show the optical density at 620nm monitored over 15 hours for an example dataset of 20 pneumococcal isolates representing different serotypes and sequence types (STs) based on the pneumococcal MLST scheme. We used the nonlinear optimisation algorithm to identify a combination of parameters with the best fit to a logistic curve based on the lowest residual sum of squares for the predicted value based on the model relative to the observed growth data (see methods). We fitted separate curves based on a subset of points for each replicate up to the end of the stationary phase or end of the experiment, whichever came first. We did this to get a better fit for the logistic curve during the growth phase, as it does not allow for negative growth, which happened after the stationary phase. We fitted the curves to infer the growth features for each of the six replicates. The averaged estimates for each replicate were then used for the downstream analyses.

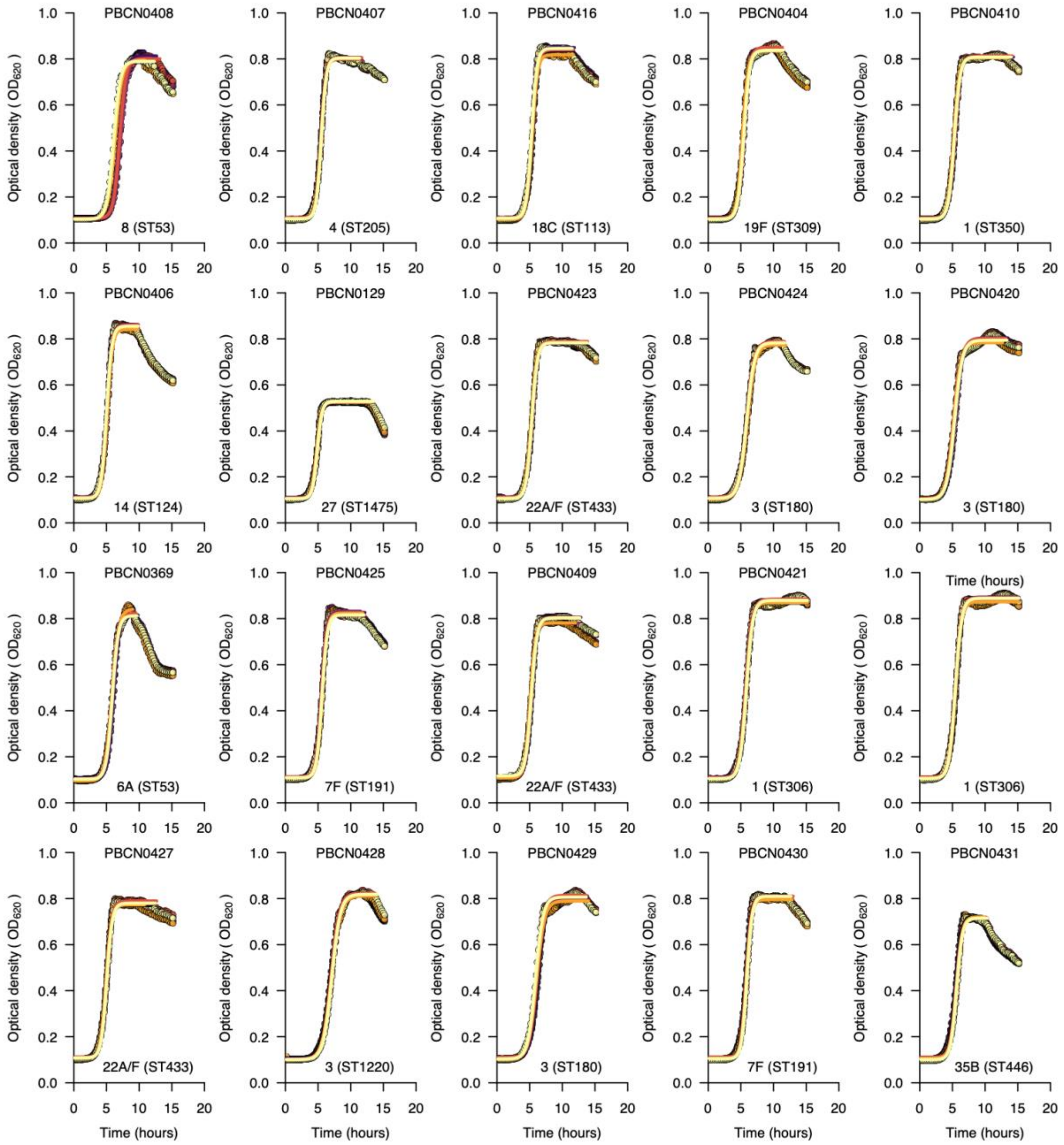


**Supplementary Fig. 1 (continued). Growth curves for selected serotypes show the *in vitro* pneumococcal growth dynamics and estimated growth features inferred by fitting a statistical model.** The growth curves above show the optical density at 620nm monitored over 15 hours for an example dataset of 20 pneumococcal isolates representing different serotypes and sequence types (STs) based on the pneumococcal MLST scheme. We used the nonlinear optimisation algorithm to identify a combination of parameters with the best fit to a logistic curve based on the lowest residual sum of squares for the predicted value based on the model relative to the observed growth data (see methods). We fitted separate curves based on a subset of points for each replicate up to the end of the stationary phase or end of the experiment, whichever came first. We did this to get a better fit for the logistic curve during the growth phase, as it does not allow for negative growth, which happened after the stationary phase. We fitted the curves to infer the growth features for each of the six replicates. The averaged estimates for each replicate were then used for the downstream analyses.

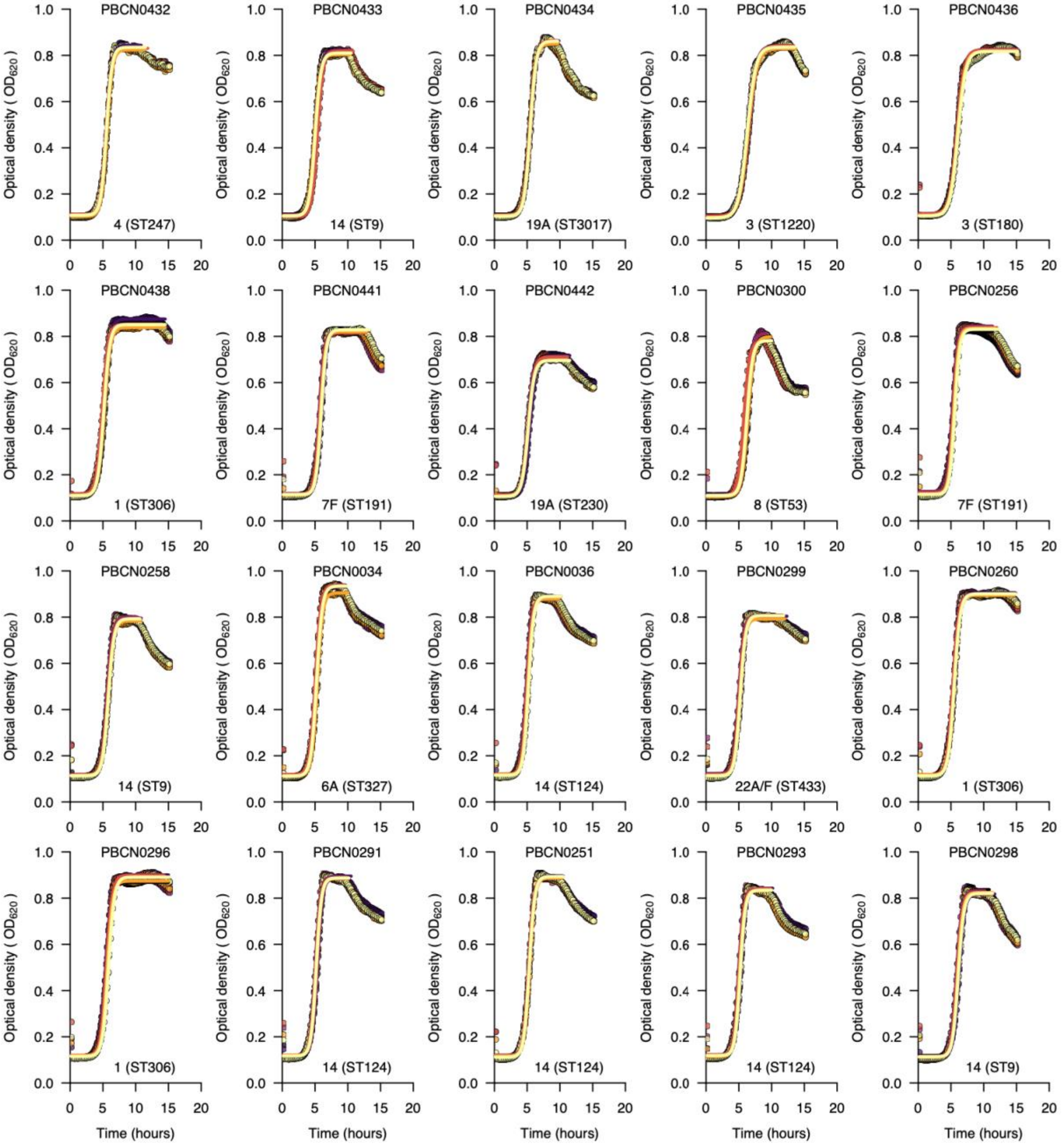


**Supplementary Fig. 1 (continued). Growth curves for selected serotypes show the *in vitro* pneumococcal growth dynamics and estimated growth features inferred by fitting a statistical model.** The growth curves above show the optical density at 620nm monitored over 15 hours for an example dataset of 20 pneumococcal isolates representing different serotypes and sequence types (STs) based on the pneumococcal MLST scheme. We used the nonlinear optimisation algorithm to identify a combination of parameters with the best fit to a logistic curve based on the lowest residual sum of squares for the predicted value based on the model relative to the observed growth data (see methods). We fitted separate curves based on a subset of points for each replicate up to the end of the stationary phase or end of the experiment, whichever came first. We did this to get a better fit for the logistic curve during the growth phase, as it does not allow for negative growth, which happened after the stationary phase. We fitted the curves to infer the growth features for each of the six replicates. The averaged estimates for each replicate were then used for the downstream analyses.

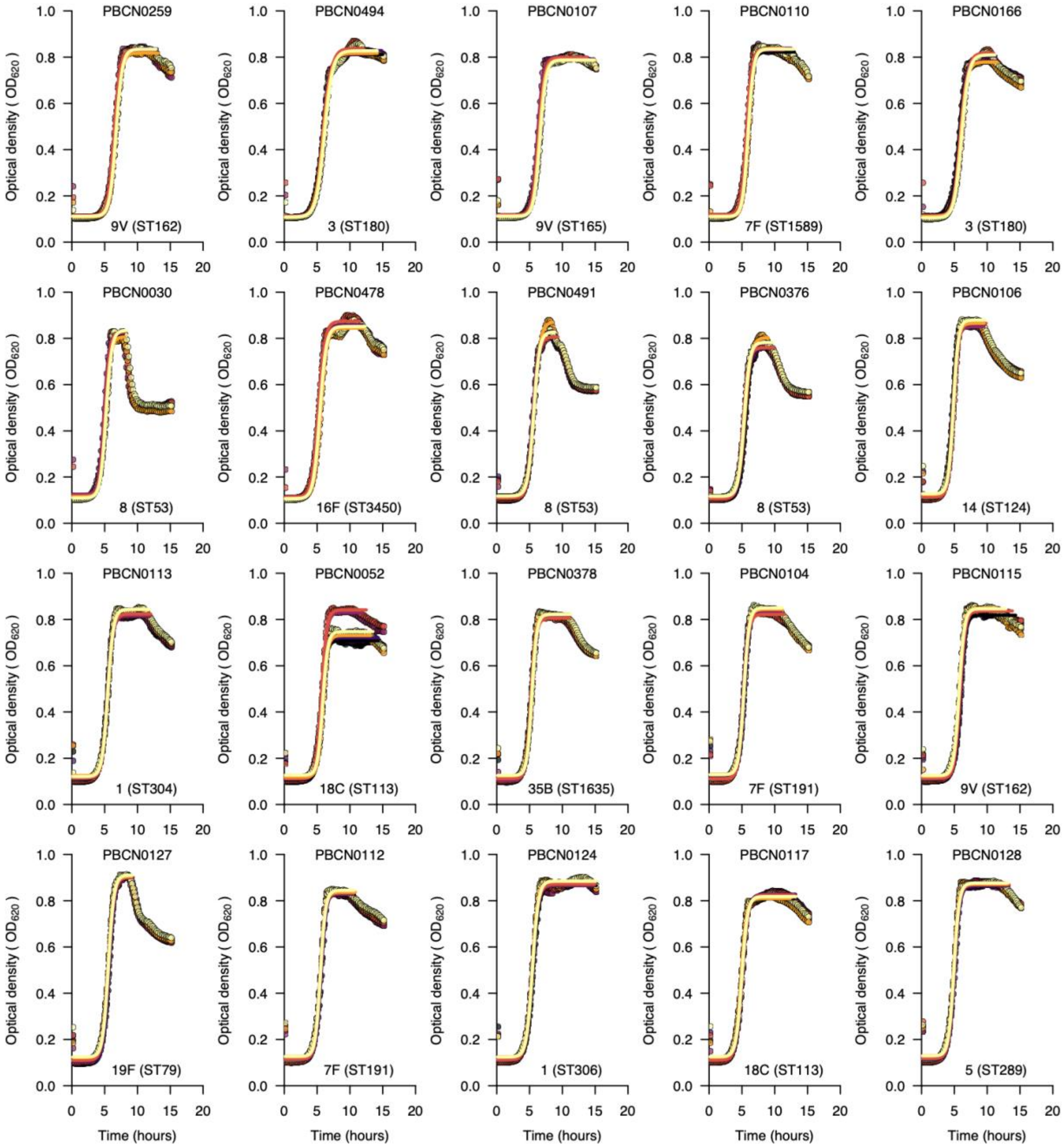


**Supplementary Fig. 1 (continued). Growth curves for selected serotypes show the *in vitro* pneumococcal growth dynamics and estimated growth features inferred by fitting a statistical model.** The growth curves above show the optical density at 620nm monitored over 15 hours for an example dataset of 20 pneumococcal isolates representing different serotypes and sequence types (STs) based on the pneumococcal MLST scheme. We used the nonlinear optimisation algorithm to identify a combination of parameters with the best fit to a logistic curve based on the lowest residual sum of squares for the predicted value based on the model relative to the observed growth data (see methods). We fitted separate curves based on a subset of points for each replicate up to the end of the stationary phase or end of the experiment, whichever came first. We did this to get a better fit for the logistic curve during the growth phase, as it does not allow for negative growth, which happened after the stationary phase. We fitted the curves to infer the growth features for each of the six replicates. The averaged estimates for each replicate were then used for the downstream analyses.

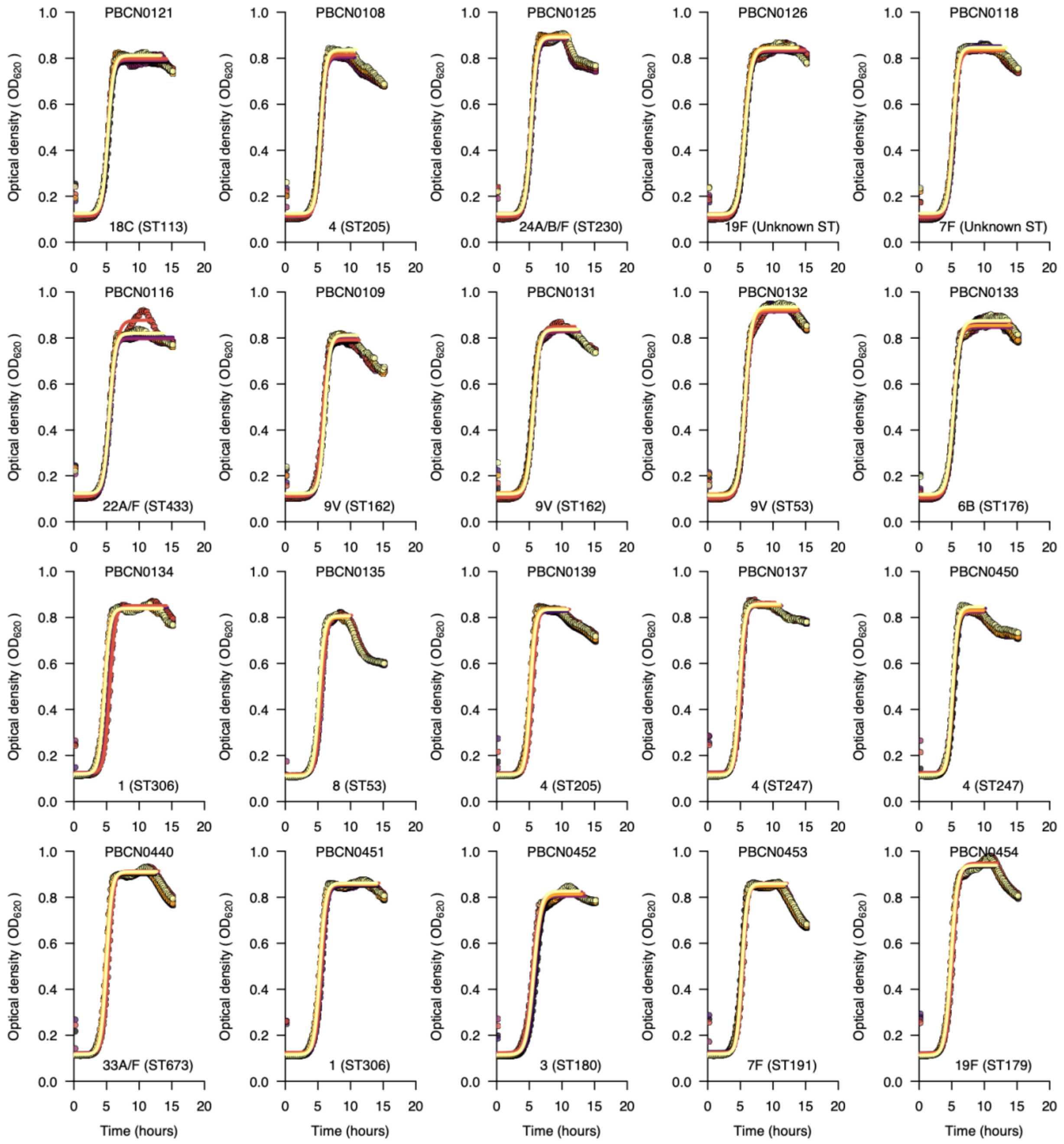


**Supplementary Fig. 1 (continued). Growth curves for selected serotypes show the *in vitro* pneumococcal growth dynamics and estimated growth features inferred by fitting a statistical model.** The growth curves above show the optical density at 620nm monitored over 15 hours for an example dataset of 20 pneumococcal isolates representing different serotypes and sequence types (STs) based on the pneumococcal MLST scheme. We used the nonlinear optimisation algorithm to identify a combination of parameters with the best fit to a logistic curve based on the lowest residual sum of squares for the predicted value based on the model relative to the observed growth data (see methods). We fitted separate curves based on a subset of points for each replicate up to the end of the stationary phase or end of the experiment, whichever came first. We did this to get a better fit for the logistic curve during the growth phase, as it does not allow for negative growth, which happened after the stationary phase. We fitted the curves to infer the growth features for each of the six replicates. The averaged estimates for each replicate were then used for the downstream analyses.

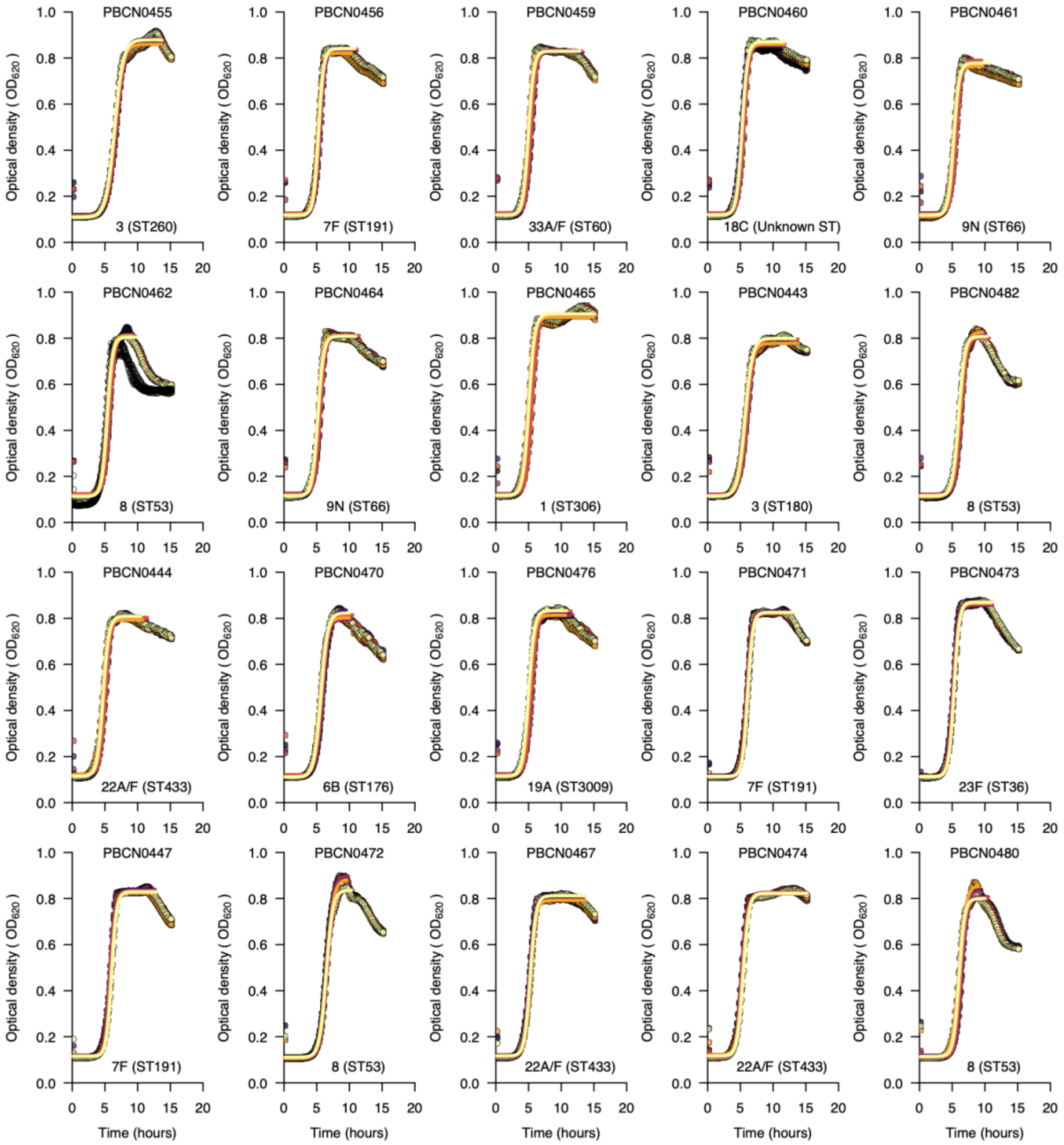


**Supplementary Fig. 1 (continued). Growth curves for selected serotypes show the *in vitro* pneumococcal growth dynamics and estimated growth features inferred by fitting a statistical model.** The growth curves above show the optical density at 620nm monitored over 15 hours for an example dataset of 20 pneumococcal isolates representing different serotypes and sequence types (STs) based on the pneumococcal MLST scheme. We used the nonlinear optimisation algorithm to identify a combination of parameters with the best fit to a logistic curve based on the lowest residual sum of squares for the predicted value based on the model relative to the observed growth data (see methods). We fitted separate curves based on a subset of points for each replicate up to the end of the stationary phase or end of the experiment, whichever came first. We did this to get a better fit for the logistic curve during the growth phase, as it does not allow for negative growth, which happened after the stationary phase. We fitted the curves to infer the growth features for each of the six replicates. The averaged estimates for each replicate were then used for the downstream analyses.

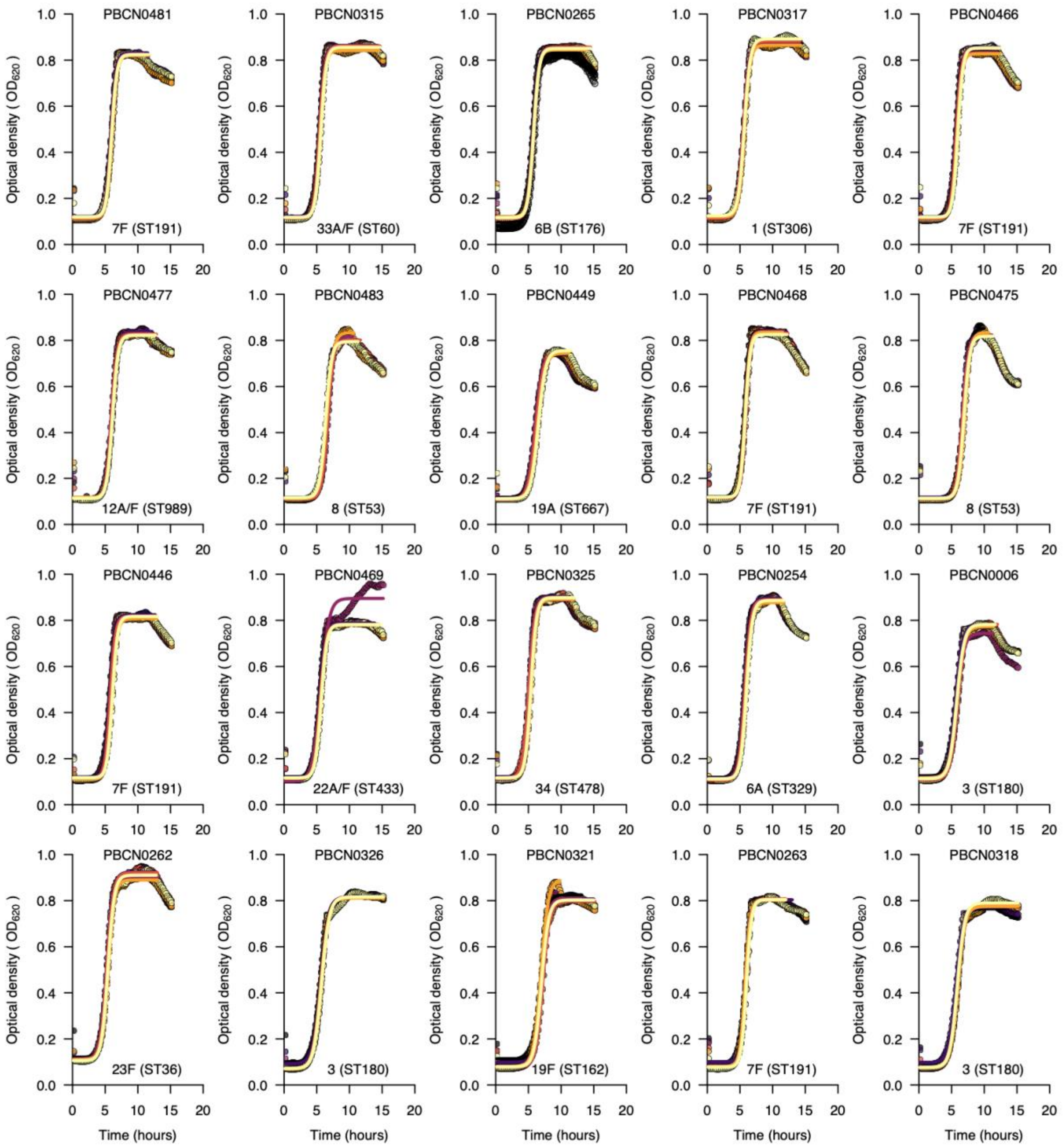


**Supplementary Fig. 1 (continued). Growth curves for selected serotypes show the *in vitro* pneumococcal growth dynamics and estimated growth features inferred by fitting a statistical model.** The growth curves above show the optical density at 620nm monitored over 15 hours for an example dataset of 20 pneumococcal isolates representing different serotypes and sequence types (STs) based on the pneumococcal MLST scheme. We used the nonlinear optimisation algorithm to identify a combination of parameters with the best fit to a logistic curve based on the lowest residual sum of squares for the predicted value based on the model relative to the observed growth data (see methods). We fitted separate curves based on a subset of points for each replicate up to the end of the stationary phase or end of the experiment, whichever came first. We did this to get a better fit for the logistic curve during the growth phase, as it does not allow for negative growth, which happened after the stationary phase. We fitted the curves to infer the growth features for each of the six replicates. The averaged estimates for each replicate were then used for the downstream analyses.

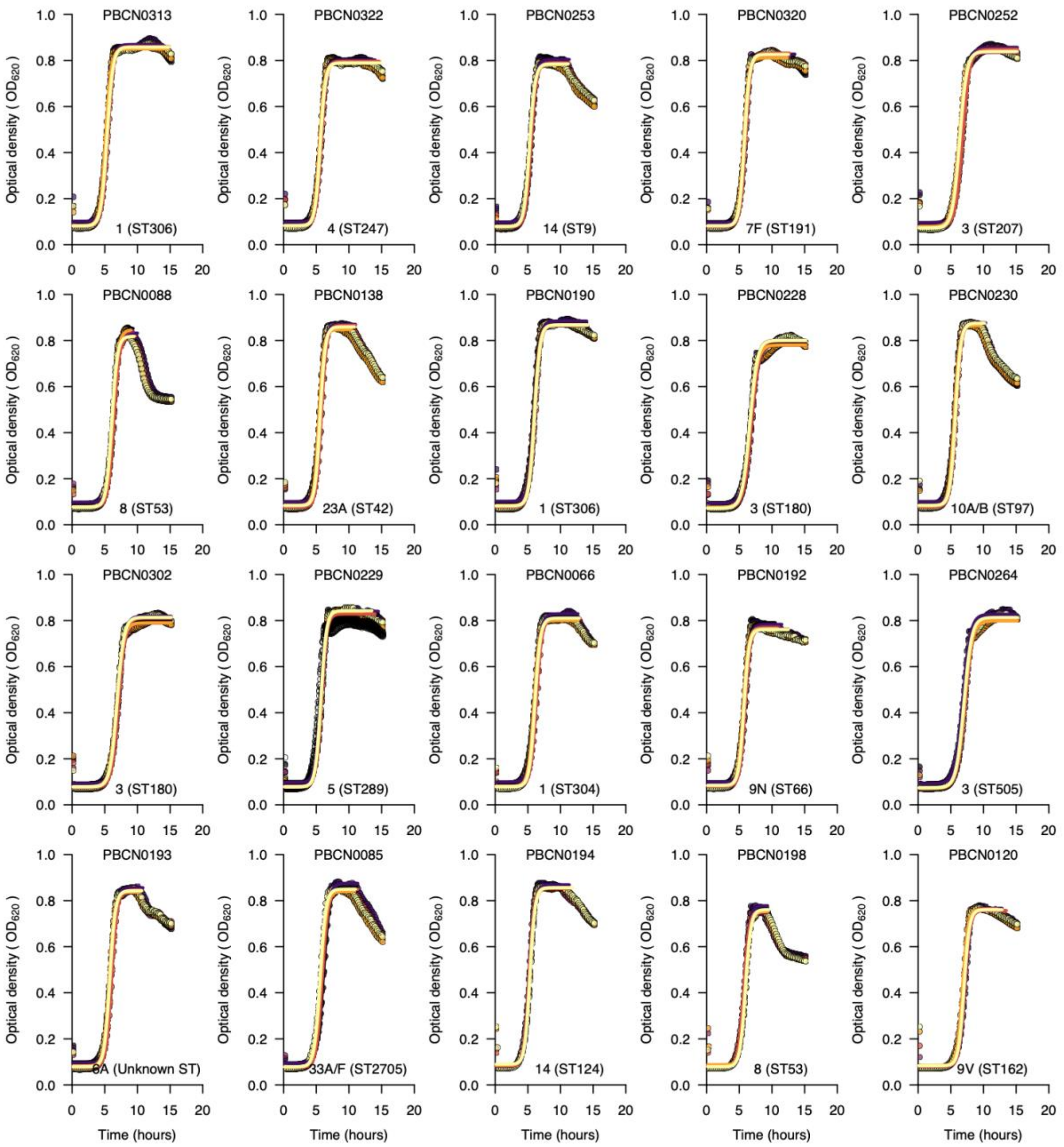


**Supplementary Fig. 1 (continued). Growth curves for selected serotypes show the *in vitro* pneumococcal growth dynamics and estimated growth features inferred by fitting a statistical model.** The growth curves above show the optical density at 620nm monitored over 15 hours for an example dataset of 20 pneumococcal isolates representing different serotypes and sequence types (STs) based on the pneumococcal MLST scheme. We used the nonlinear optimisation algorithm to identify a combination of parameters with the best fit to a logistic curve based on the lowest residual sum of squares for the predicted value based on the model relative to the observed growth data (see methods). We fitted separate curves based on a subset of points for each replicate up to the end of the stationary phase or end of the experiment, whichever came first. We did this to get a better fit for the logistic curve during the growth phase, as it does not allow for negative growth, which happened after the stationary phase. We fitted the curves to infer the growth features for each of the six replicates. The averaged estimates for each replicate were then used for the downstream analyses.

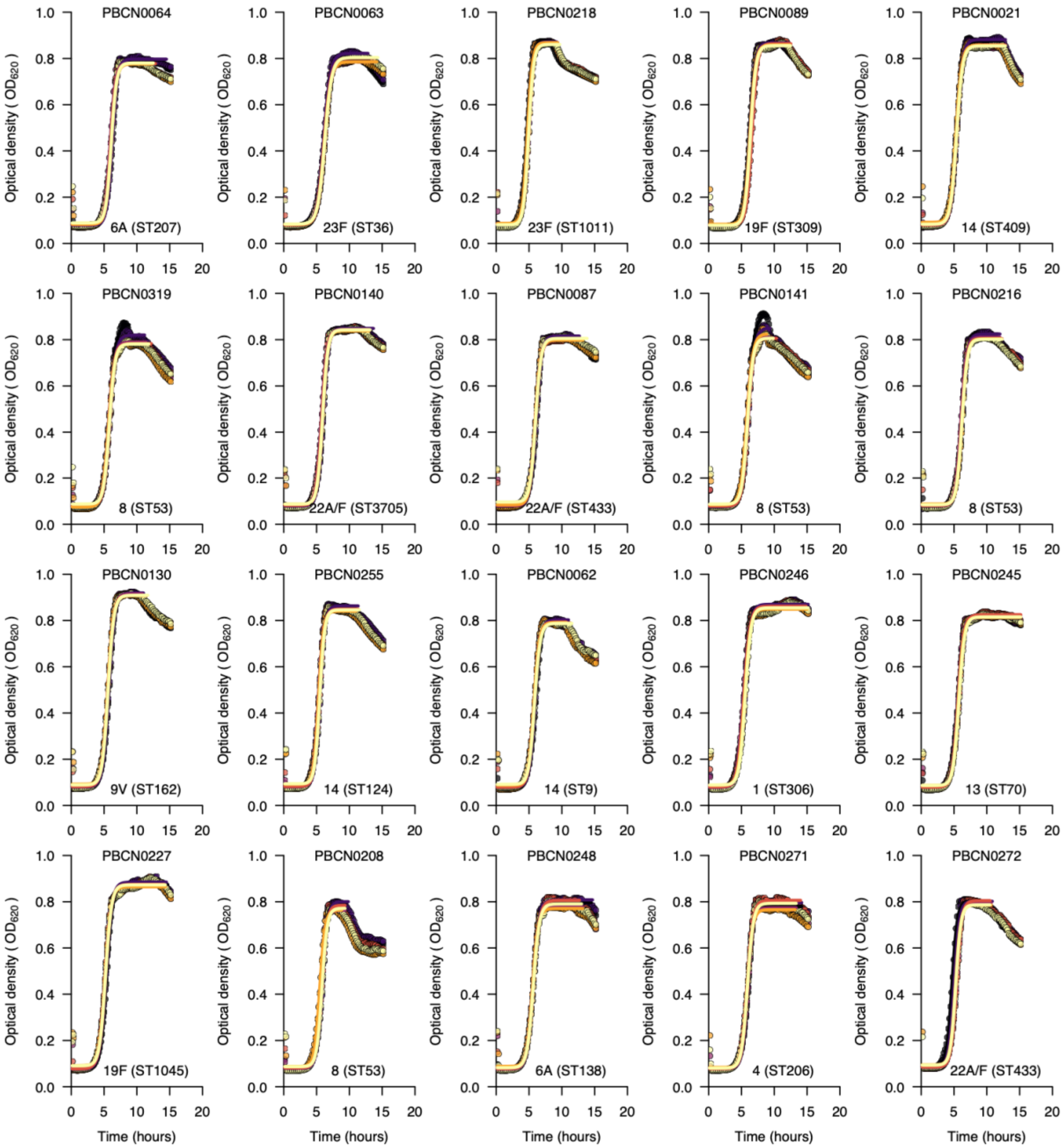


**Supplementary Fig. 1 (continued). Growth curves for selected serotypes show the *in vitro* pneumococcal growth dynamics and estimated growth features inferred by fitting a statistical model.** The growth curves above show the optical density at 620nm monitored over 15 hours for an example dataset of 20 pneumococcal isolates representing different serotypes and sequence types (STs) based on the pneumococcal MLST scheme. We used the nonlinear optimisation algorithm to identify a combination of parameters with the best fit to a logistic curve based on the lowest residual sum of squares for the predicted value based on the model relative to the observed growth data (see methods). We fitted separate curves based on a subset of points for each replicate up to the end of the stationary phase or end of the experiment, whichever came first. We did this to get a better fit for the logistic curve during the growth phase, as it does not allow for negative growth, which happened after the stationary phase. We fitted the curves to infer the growth features for each of the six replicates. The averaged estimates for each replicate were then used for the downstream analyses.

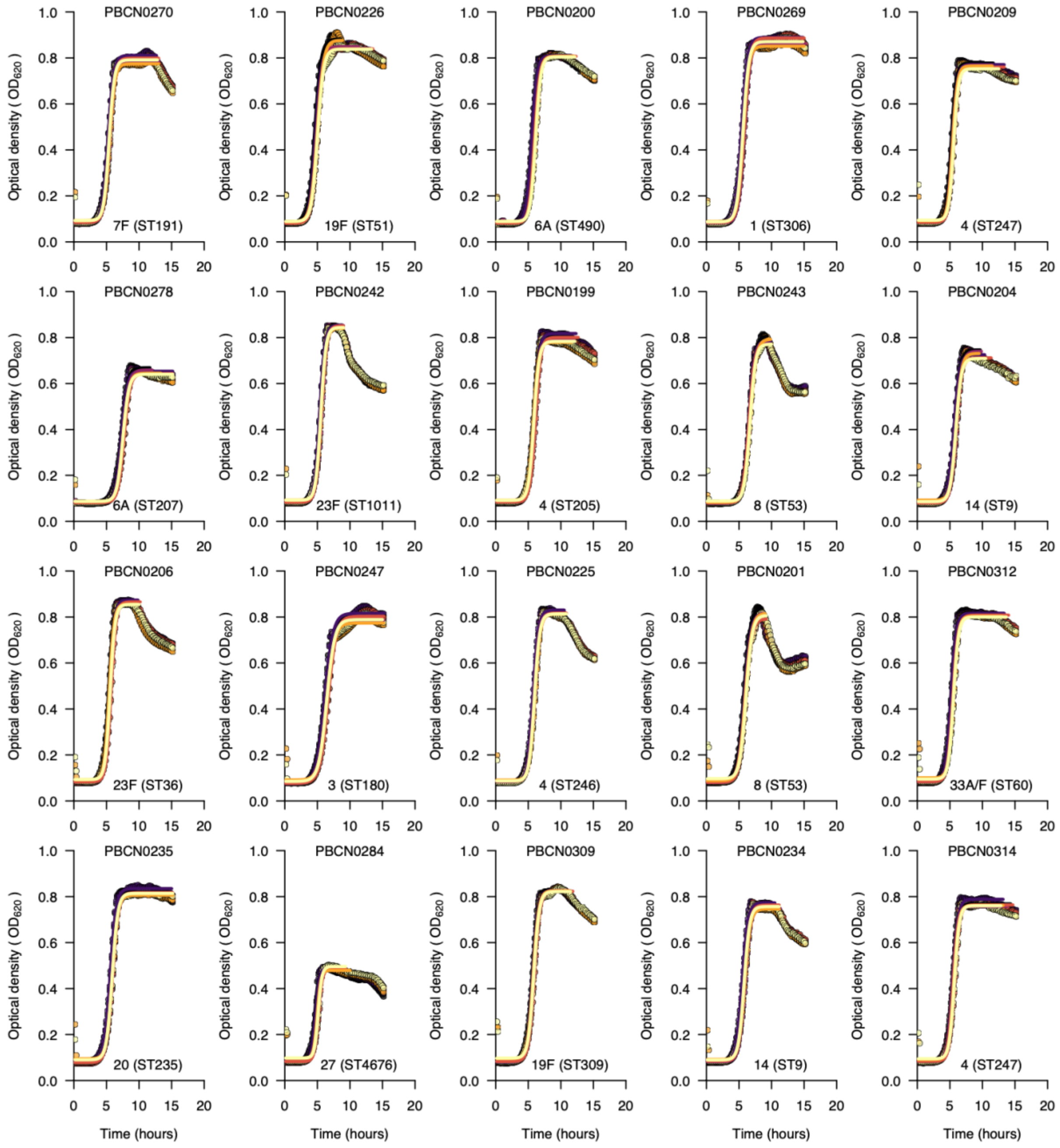


**Supplementary Fig. 1 (continued). Growth curves for selected serotypes show the *in vitro* pneumococcal growth dynamics and estimated growth features inferred by fitting a statistical model.** The growth curves above show the optical density at 620nm monitored over 15 hours for an example dataset of 20 pneumococcal isolates representing different serotypes and sequence types (STs) based on the pneumococcal MLST scheme. We used the nonlinear optimisation algorithm to identify a combination of parameters with the best fit to a logistic curve based on the lowest residual sum of squares for the predicted value based on the model relative to the observed growth data (see methods). We fitted separate curves based on a subset of points for each replicate up to the end of the stationary phase or end of the experiment, whichever came first. We did this to get a better fit for the logistic curve during the growth phase, as it does not allow for negative growth, which happened after the stationary phase. We fitted the curves to infer the growth features for each of the six replicates. The averaged estimates for each replicate were then used for the downstream analyses.

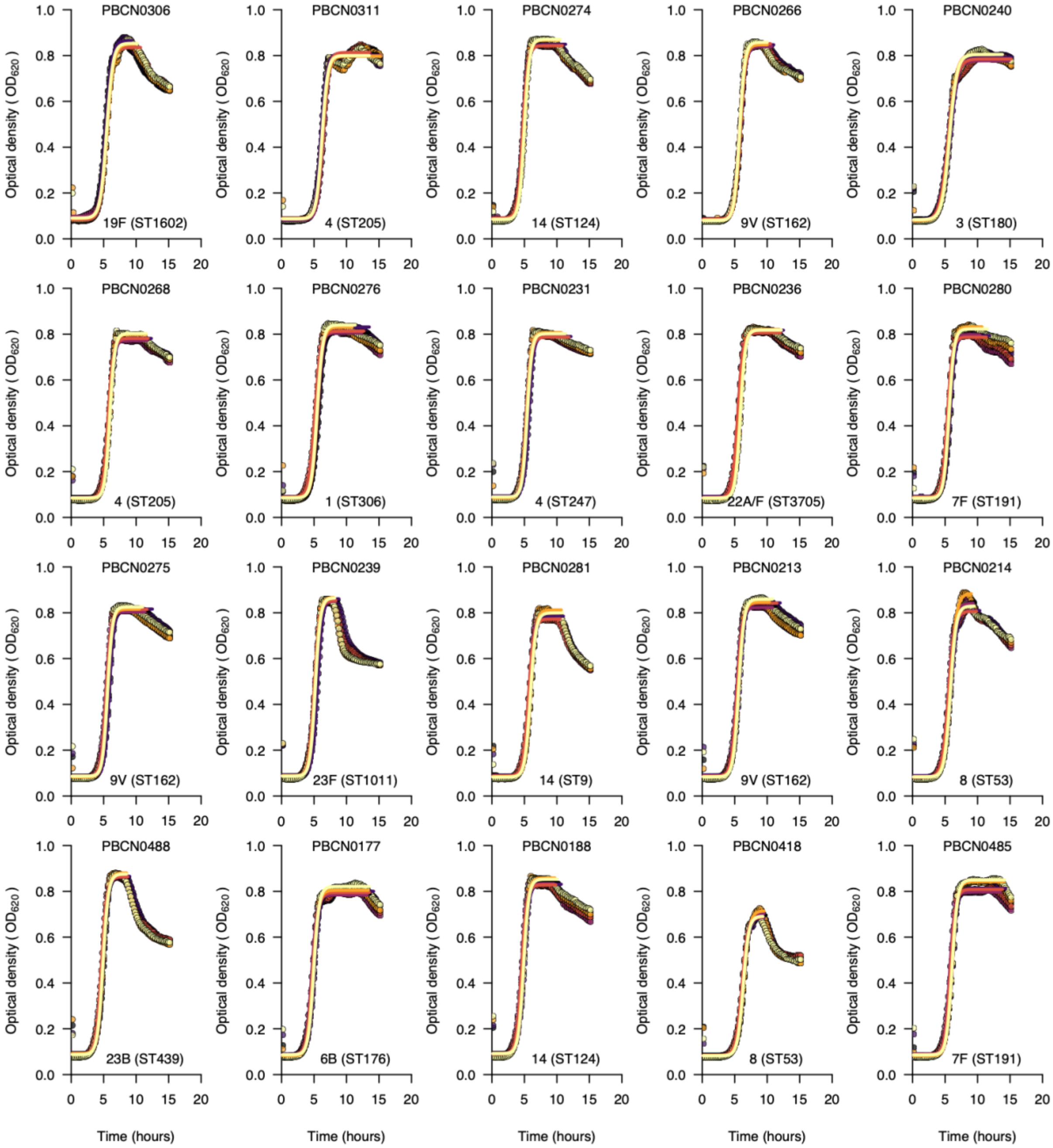


**Supplementary Fig. 1 (continued). Growth curves for selected serotypes show the *in vitro* pneumococcal growth dynamics and estimated growth features inferred by fitting a statistical model.** The growth curves above show the optical density at 620nm monitored over 15 hours for an example dataset of 20 pneumococcal isolates representing different serotypes and sequence types (STs) based on the pneumococcal MLST scheme. We used the nonlinear optimisation algorithm to identify a combination of parameters with the best fit to a logistic curve based on the lowest residual sum of squares for the predicted value based on the model relative to the observed growth data (see methods). We fitted separate curves based on a subset of points for each replicate up to the end of the stationary phase or end of the experiment, whichever came first. We did this to get a better fit for the logistic curve during the growth phase, as it does not allow for negative growth, which happened after the stationary phase. We fitted the curves to infer the growth features for each of the six replicates. The averaged estimates for each replicate were then used for the downstream analyses.

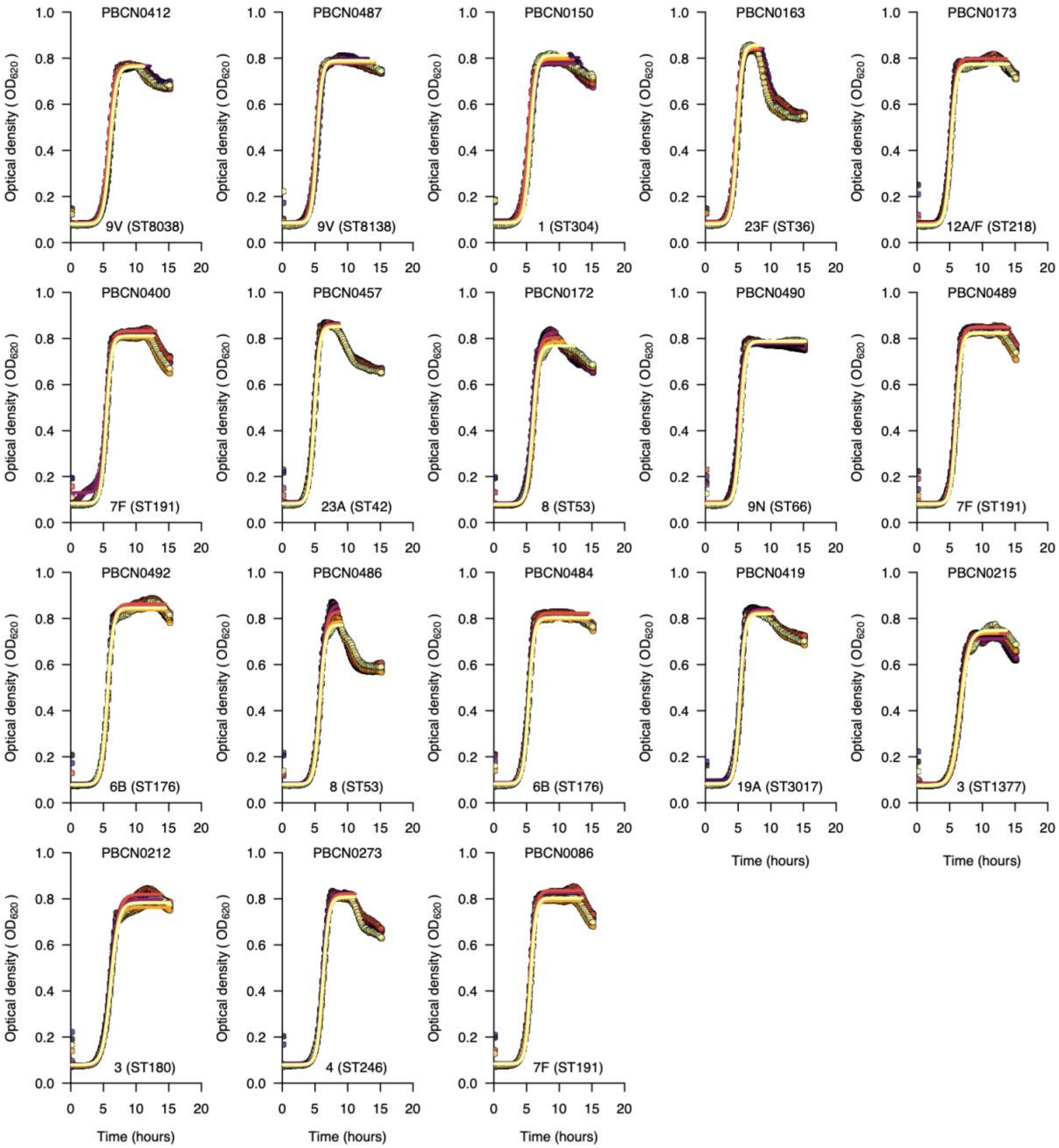


**Supplementary Fig. 1 (continued). Growth curves for selected serotypes show the *in vitro* pneumococcal growth dynamics and estimated growth features inferred by fitting a statistical model.** The growth curves above show the optical density at 620nm monitored over 15 hours for an example dataset of 20 pneumococcal isolates representing different serotypes and sequence types (STs) based on the pneumococcal MLST scheme. We used the nonlinear optimisation algorithm to identify a combination of parameters with the best fit to a logistic curve based on the lowest residual sum of squares for the predicted value based on the model relative to the observed growth data (see methods). We fitted separate curves based on a subset of points for each replicate up to the end of the stationary phase or end of the experiment, whichever came first. We did this to get a better fit for the logistic curve during the growth phase, as it does not allow for negative growth, which happened after the stationary phase. We fitted the curves to infer the growth features for each of the six replicates. The averaged estimates for each replicate were then used for the downstream analyses.

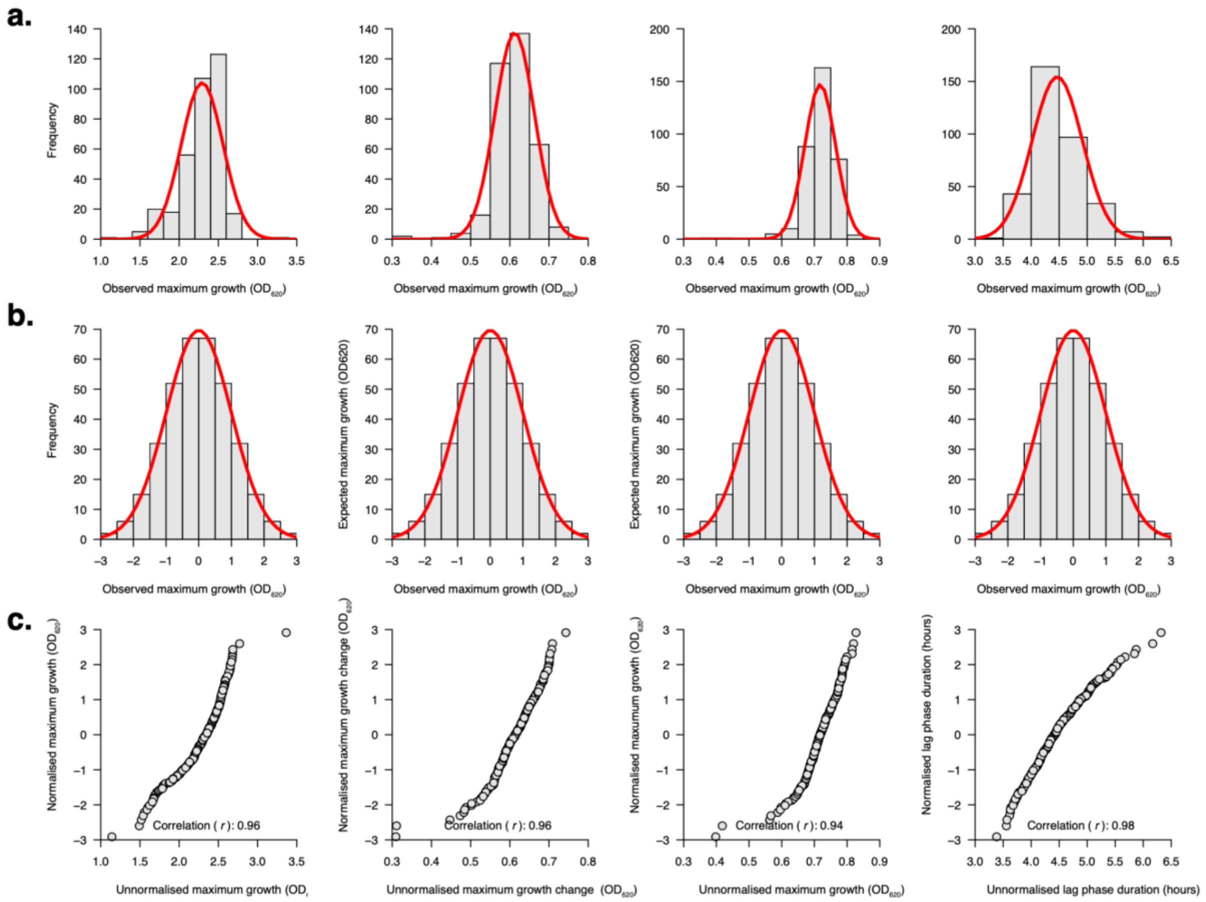


**Supplementary Fig. 2. Correlation between the normalised and unnormalised growth parameters.** The histograms show the distribution of the estimated growth features: average growth rate, maximum growth change, maximum growth density, and lag phase duration (from left to right) when **(a)** unnormalised and **(b)** normalised using rank-based inverse normal transformation (see methods). The pneumococcal growth parameters were normalised using rank-based inverse normal transformation (https://cran.r-project.org/web/packages/RNOmni/) prior to the GWAS analysis. **(c)** Scatterplots showing correlation between the unnormalised and normalised growth features. Correlation was measured using the Pearson correlation coefficient.

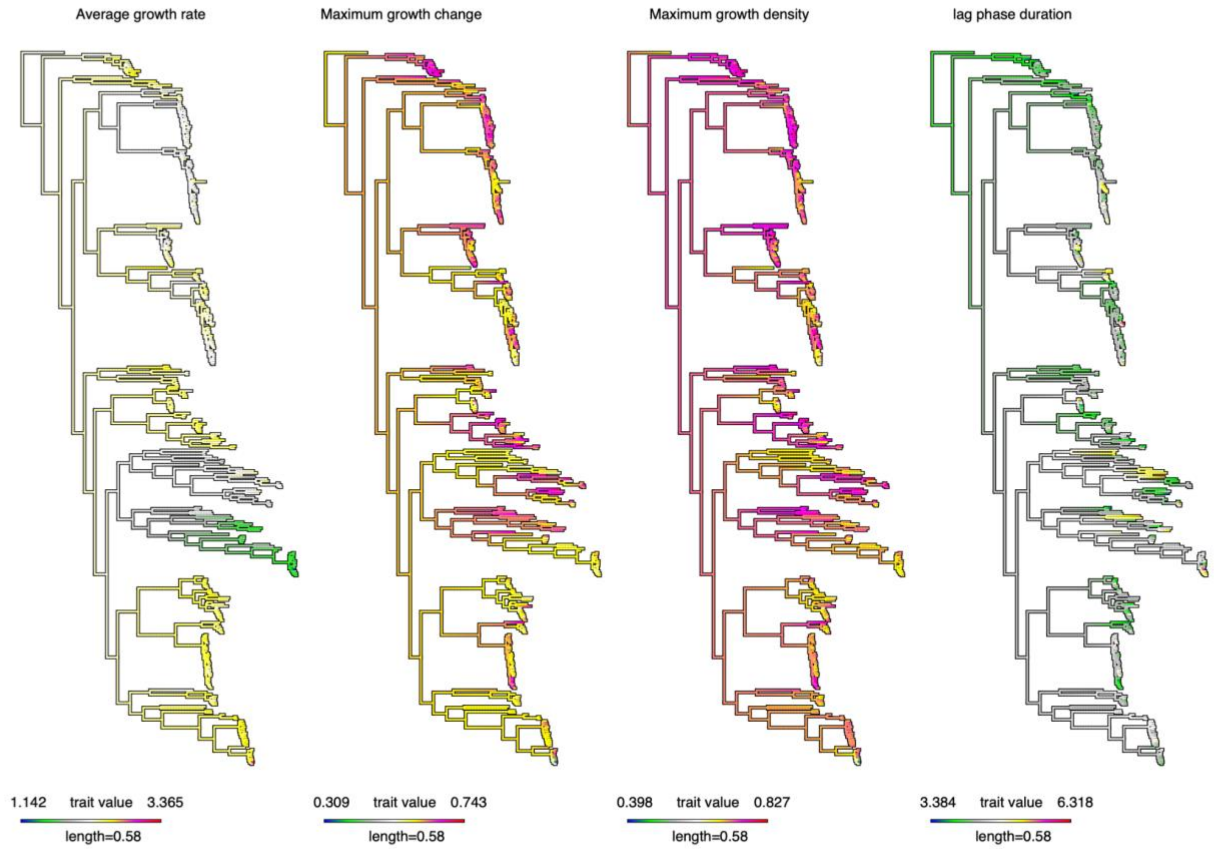


**Supplementary Fig. 3. Ancestral reconstruction reveals a strong correlation between the estimated *in vitro* pneumococcal growth features and the whole-genome phylogeny of the isolates.** The evolution of the unnormalised estimated continuous growth features, namely average growth rate, maximum growth change, maximum growth density, and lag phase duration, was assessed by estimating the states of the internal nodes of the maximum likelihood phylogenetic tree of the isolates using a maximum likelihood ancestral reconstruction approach. The length of the scale bar shows the number of nucleotide substitutions per site, while the colours represent the lowest and highest unnormalised growth features.
